# Impairment but not abolishment of express saccades after unilateral- or bilateral cryogenic FEF inactivation

**DOI:** 10.1101/583823

**Authors:** Suryadeep Dash, Tyler R. Peel, Stephen G. Lomber, Brian D. Corneil

**Affiliations:** Department of Physiology & Pharmacology, University of Western Ontario, London, ON, Canada; Robarts Research Institute, University of Western Ontario, London, ON, Canada; Graduate Program in Neuroscience, University of Western Ontario, London, ON, Canada; Department of Psychology, University of Western Ontario, London, ON, Canada

**Keywords:** Oculomotor system, express saccades, frontal eye fields, motor preparation

## Abstract

Express saccades (ESs) are a manifestation of a visual grasp reflex triggered when visual information arrives in the intermediate layers of the superior colliculus (SCi), which in turn orchestrates the lower level brainstem saccade generator to evoke a saccade with a very short latency (∼100ms). A prominent theory regarding express saccades generation is that they are facilitated by preparatory signals, presumably from cortical areas, which prime the SCi prior to the arrival of visual information. Here, we test this theory by reversibly inactivating a key cortical input to the SCi, the frontal eye fields (FEF), while monkeys perform an oculomotor task that promotes ES generation. Across three tasks with a different combination of potential target locations and uni- or bilateral FEF inactivation, we found a spared ability for monkeys to generate ESs, despite decreases in ES frequency during FEF inactivation. This result is consistent with the FEF having a facilitatory but not critical role in ES generation, likely because other cortical areas compensate for the loss of preparatory input to the SCi. However, we did find decreases in the accuracy and peak velocity of ESs generated during FEF inactivation, which argues for an influence of the FEF on the saccadic burst generator even during ESs. Overall, our results shed further light on the role of the FEF in the shortest-latency visually-guided eye movements.

**New & Noteworthy:** Express saccades (ESs) are the shortest-latency visually-guided saccade. The frontal eye fields (FEF) is thought to promote ES by establishing the necessary preconditions in the superior colliculus. Here, by reversibly inactivate the FEF either unilaterally or bilaterally, we support this view by showing that the FEF plays an assistive but not critical role in ES generation. We also found that FEF inactivation lowered ES peak velocity, emphasizing a contribution of the FEF to ES kinematics.

## Introduction

Express saccades (ESs) are the shortest-latency visually-guided saccades, with reaction times (RTs) approaching the sensori-motor conduction delays between the retina and extra-ocular muscles (Fischer and Boch 1983; Fischer and Ramsperger 1984). Express saccades are essentially a low-level visual grasp reflex that, somewhat paradoxically, is potentiated by high-level preparation about the location, valence, or timing of an upcoming visual target (Paré and Munoz 1996; Schiller et al. 2004). The interaction between a low-level reflex and high-level preparation is seen in the activity profiles within the intermediate superior colliculus (SCi), a structure whose integrity is essential for ES generation (Schiller et al. 1987). Before regular (or non-express) latency saccades, a subset of *visuomotor* neurons within the SCi emit a ‘visual’ burst of action potentials shortly after visual target presentation, as well as a second ‘motor’ burst of action potentials shortly before the onset of a target-directed saccade (Dorris et al. 1997; Edelman and Keller 1996; Sparks et al. 2000). Prior to ESs, visuomotor neurons emit only a single burst of action potentials; effectively the visual and motor events become a singular event linked to both target and saccade onset (Dorris et al. 1997; Edelman and Keller 1996; Sparks et al. 2000). Within the SCi, a neural correlate of high-level preparation that potentiates ES generation is the level of low-frequency activity on SCi visuomotor neurons attained just before the arrival of the visual burst of activity. Although the exact details determining saccade triggering within the SCi and downstream brainstem burst generation remain to be determined, greater levels of low-frequency activity within the SCi are thought to bring the system closer to saccade threshold, increasing the probability of ES generation when the visually-related burst of activity arrives in the SCi (Dash et al. 2018; Dorris et al. 1997; Krauzlis 2003; Rezvani and Corneil 2008).

There are many cortical and subcortical areas that exhibit preparatory-related activity and project directly to the SCi, and hence could provide the high-level signals needed to potentiate ES generation (Ashmore and Sommer 2013; Chen et al. 2013; Hikosaka et al. 1989; Johnston et al. 2014; Kim and Lee 2017; Ohmae et al. 2017; Wurtz and Hikosaka 1986). The frontal eye fields (FEF) in particular is an important source of top-down signals conveyed to the SCi (Sommer and Wurtz 2000; 2001; Wurtz et al. 2001), and many FEF neurons also display increased levels of low-frequency preparatory activity that correlates with increased ES probability (Dias and Bruce 1994; Everling and Munoz 2000). Somewhat surprisingly in light of these findings, the influence of temporary or permanent lesions of the FEF on ES occurrence is quite varied. On one hand, monkeys with permanent unilateral lesion of the FEF are still capable of generating contralesional ESs after a short period of recovery (Schiller et al. 1987), and both ipsilesionally-and contralesionally-directed ESs are also spared, and sometimes potentiated, by unilateral damage to the FEF in human patients (Braun et al. 1992; Guitton et al. 1985; Rivaud et al. 1994). In contrast, we have recently shown in monkeys that low-frequency preparatory activity and visual-and saccade-related activity in the ipsilesional SCi decrease with reversible cryogenic inactivation of a large-volume of the unilateral FEF (Dash et al. 2018; Peel et al. 2017). The study by Dash and colleagues (2018) also showed that the profile of SCi activity prior to ESs generation is largely unchanged during FEF inactivation, providing that low-frequency preparatory activity in the SCi reaches sufficient levels presumably due to inputs from other non-FEF sources (Dash et al. 2018). Thus, at least when long-term compensation is not involved, the FEF appears to play a facilitatory but not critical role for ES generation.

The overall goal of the current manuscript is to further the understanding of the FEF’s role in ES generation, here by examining ES probability, accuracy, and peak velocity in monkey before, during, and after either unilateral or bilateral cryogenic inactivation of the FEF. To our knowledge, there has been no study of the effect of bilateral FEF inactivation on ES generation, despite recent evidence showing a role for the FEF in bilateral saccade generation (Crapse and Sommer 2009; Kunimatsu et al. 2015; Peel et al. 2014). Further, as SCi activity was not recorded in the current study, we were not limited to the two-target configuration used in the study by Dash and colleagues (2018), and here introduced multiple potential target locations either unilaterally or bilaterally during FEF inactivation. Doing so allowed us to examine how uni- or bilateral FEF inactivation interacts with different potential target locations to influence ES generation. Finally, we also examined ES accuracy and peak velocity during FEF inactivation to gain further understanding into the FEF’s potential influence on the movement parameters of ES. We found that while ES frequency decreased after either uni- or bilateral FEF inactivation, the ability for the NHPs to generate ESs was never completely abolished. When generated, ESs made during FEF inactivation tended to be slower and more inaccurate. Overall, our study reaffirms the potentiating but not critical role for the FEF in ESs, and highlights the FEF’s continuing influence on saccadic kinematics regardless of reaction time.

## Method

### Subjects and surgical procedures

Two male rhesus monkeys (*Macaca mulatta*, DZ, and OZ weighing 9.8, and 8.6 kg respectively) were prepared for head immobilization and cryogenic inactivation of FEF. All training, surgical, and experimental procedures were in accordance with the Canadian Council on Animal Care policy on the use of laboratory animals and approved by the Animal Use Subcommittee of the University of Western Ontario Council on Animal Care. Surgical procedures describing drug regimes, post-surgical care and implantation of head post and cryoloops, cyroloop dimensions, and estimated volume of inactivation for the monkeys used in this study can be found elsewhere (Peel et al. 2017; Peel et al. 2016; Peel et al. 2014). Briefly, each monkey was implanted bilaterally with two stainless steel cryoloops in the inferior and superior aspects of the arcuate sulcus [inferior arm (IA), superior arm (SA)]. In this study we have only used IA cooling in both monkeys as this increases trial yield during cooling, and produces ∼70% of the SRT deficits caused by combined unilateral cooling of the IA and SA (Peel et al. 2014). We have estimated that cooling an individual IA cryoloop inactivates a 90 mm^3^ volume of tissue (Peel et al. 2017; Peel et al. 2016).

### Experimental procedures

Monkeys were seated in a custom-made primate chair with their head immobilized, facing a rectilinear grid of >500 red LEDs covering ± 35° of the horizontal and vertical visual field. Eye movements were recorded using a single, chair-mounted eye tracker (EyeLink II, resolution = 0.05°, sampling rate 500Hz). All experiments were conducted in a dark, sound-attenuated room. The behavioral tasks were controlled by customized real-time LabView programs running on a PXI controller (National Instruments) at a rate of 1 kHz.

An experimental dataset consisted of pre-cooling, peri-cooling, and post-cooling sessions, with 80-150 correct trials within each session (i.e., between 240-450 trials total). After the pre-cooling session, the cooling pumps were turned on allowing the flow of chilled methanol through the lumen of the cryoloops. The peri-cooling session was initiated when cryoloop temperature reached and stayed stable at 3°C. Once sufficient data were collected for the peri-cooling session, cooling pumps were turned off, which allowed the cryoloop temperature to rapidly return towards body temperature. When cryoloop temperature exceeded 35°C, the post-cooling session was initiated.

### Behavioral task

Monkeys performed a visually-guided gap saccade task to look to peripheral targets placed on the horizontal meridian. The disengagement of active visual fixation during this task promotes ESs, but fixation disengagement is not a pre-requisite for ES generation (Munoz et al. 2000). Monkeys fixated the central fixation LED for 750-1000ms until it disappeared, which was followed by a period of 200ms (the gap interval) during which the monkeys maintained central fixation. A peripheral visual target was then presented for 150ms (the locations of potential peripheral targets are described below). To receive a fluid reward, monkeys had to initiate a saccade to the peripheral target within 500 ms, and land within a circular spatial window with a diameter of 60% of target’s visual eccentricity. The large spatial tolerance window was based on the expected decreases in saccade amplitude and accuracy following FEF inactivation (Dias and Segraves 1999; Peel et al. 2014; Sommer and Tehovnik 1997).

All behavioral analyses were carried out using customized MATLAB programs (MATLAB, The MathsWorks Inc., MA). Eye position traces were filtered using a 3rd order low pass butterworth filter and differentiated to produce eye velocity. Eye velocity was used to determine the onset and offset times of saccades with a velocity criterion of 30°/s and the maximum instantaneous velocity between saccade onset and offset was deemed as peak velocity. We analyzed the first saccade following peripheral target presentation. The eye position at saccade offset was used to calculate the mean saccade amplitude (horizontal and vertical) and end-point scatter, which was calculated as the mean angular distance between the displacements of mean and individual saccade end points from the central fixation position. Visual inspection of the data off-line confirmed the validity of automatic marking. Across our entire dataset, we observed accurate saccades to visual targets with SRTs as low as 50 ms, even in blocks where multiple potential target locations were possible to the left or right. Very few saccades were generated with SRTs < 50ms, and these were classified as anticipatory and discarded.

### Experimental design and statistical analysis

We performed 3 different experiments, with each experiment consisting of a unique combination of target locations and cooling configurations. In experiment 1 (8 or 9 sessions in monkeys DZ or OZ, respectively), unilateral inactivation of left FEF was paired with contralateral target presentation (3 potential locations). In experiment 2 (18 or 13 sessions in monkeys DZ or OZ, respectively), unilateral inactivation of the left FEF was paired with bilateral target presentation (6 potential locations, 3 per side). In experiment 3 (6 or 5 sessions in DZ or OZ, respectively), bilateral FEF inactivation was paired with bilateral target presentation (same targets as experiment 2). In all experiments, potential targets were placed at different eccentricities along the horizontal meridian (Monkey DZ: 10, 14 and 20; Monkey OZ: 6, 12 and 20).

Consistent with previous studies (Dash et al. 2018; Paré and Munoz 1996; Sparks et al. 2000), we defined express saccades as saccades with SRTs between 50ms and 120ms. We report the comparison of data collected from pre-and post-cooling sessions (termed FEFwarm) to data collected from the peri-cooling sessions (FEFcool); similar results were obtained if data from peri-cooling sessions was compared to data from pre-and post-cooling sessions separately. Data collected from different sessions in the same Experiment were pooled together. In this study, we compared the effects of FEF inactivation on SRT, the proportion of express saccades (ES), peak velocity, saccade targeting error (mean vectorial distance between target amplitude and individual saccade endpoint), and saccade end-point scatter (i.e, the vectorial difference between mean saccade endpoint and individual saccade endpoint). Since many of these measures differed for different target locations (see results), we performed separate 3-way ANOVAs to compare how a given measure (e.g., SRT) changed with amplitude (small, medium and large), experimental configuration (Exp1, Exp2 and Exp3) and inactivation (FEFwarm and FEFcool) as the three 3 factors. Posthoc analyses comparing the subgroups were performed using Tukey-kramer test corrected for multiple comparisons. Finally, since FEF inactivation changes saccade metrics, comparisons of peak velocity were performed with a Wilcoxon sign rank task (α < 0.05) after extracting pairs of saccades, one each from the FEFwarm and FEFcool dataset, whose horizontal and vertical components differed by < 0.5 deg.

## Results

### Express saccades persist during bilateral FEF inactivation

We studied the effects of reversible unilateral or bilateral FEF inactivation on saccade reaction times (SRT), frequency of express saccades (ES), and metrics and kinematics of both ES and regular-latency saccades during a gap-saccade task. Figure 1 shows exemplary and group results for the configuration expected to yield the maximum effect, when the FEF was inactivated bilaterally (i.e., Experiment 3). In this figure, red traces and histograms represent data collected when the FEF was warm, and blue traces and histograms represent data collected when the FEF was inactivated.

**Figure 1:**
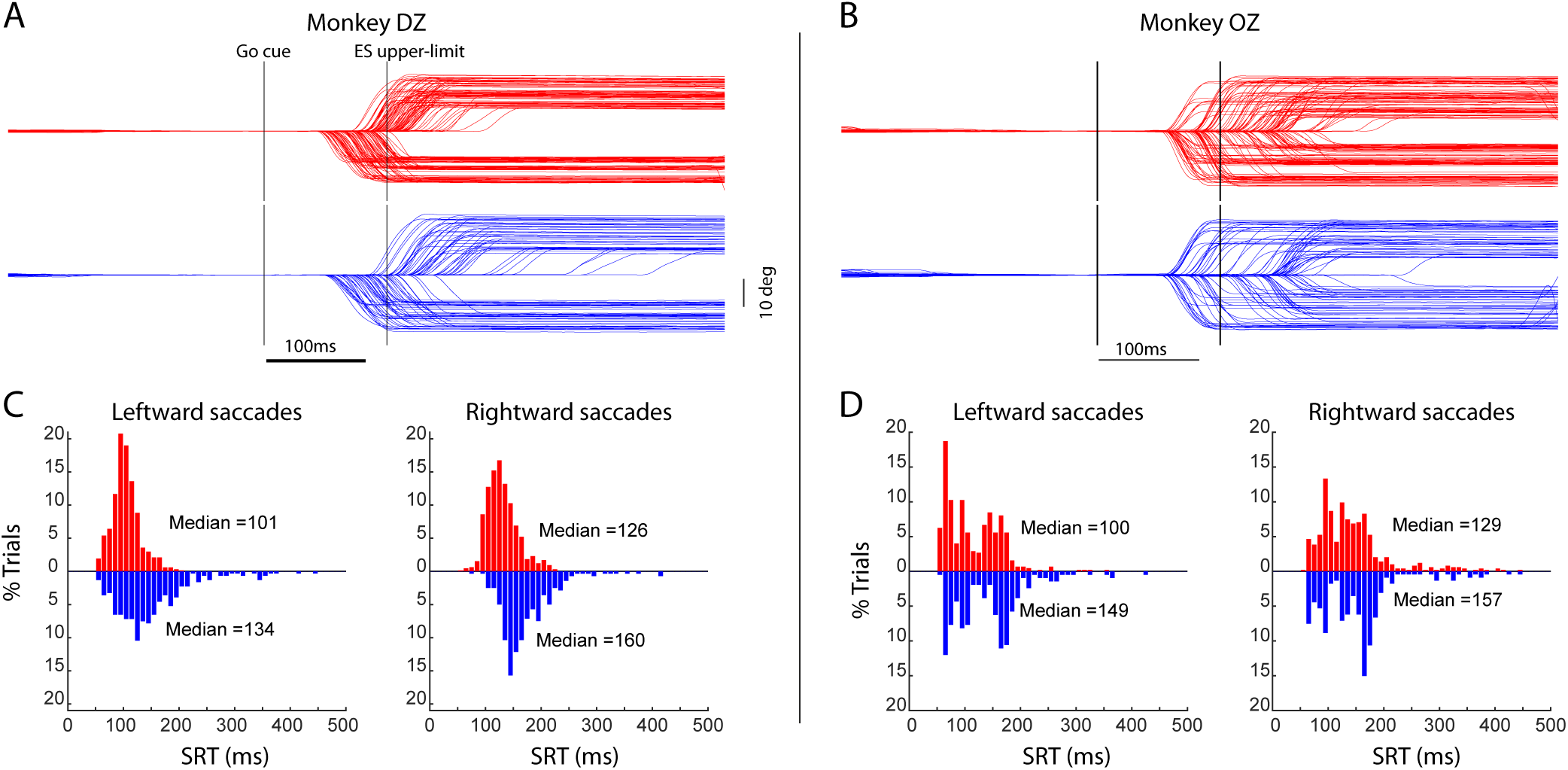
Effect of FEF inactivation on SRT. **A & B:** Example session with bilateral FEF inactivation in monkey DZ and monkey OZ, respectively. Rightward and leftward saccades are represented by positive and negative changes in eye position, respectively. Red traces represent FEFwarm trials and blue traces represents FEFcool trials. **C & D:** SRT histograms for FEFwarm (red) and FEFcool (blue) trials across all the sessions of bilateral FEF inactivation in monkey DZ and monkey OZ, respectively. All target amplitudes are pooled together in these histograms.

Recall in Experiment 3 that there were six possible target locations (3 in each direction). Despite this uncertainty, both animals generated very short-RT saccades in the ES range when the FEF was not inactivated. Close inspection of these trials reveals that even the shortest-RT movements landed near the flashed visual target, as shown by the banding of eye movement traces in each direction, corresponding to the 3 target amplitudes. Monkey DZ also exhibited a prominent SRT asymmetry, with leftward saccades having shorter SRTs than rightward saccades. Further, and as is particularly clear in the SRT histograms of data collected across all sessions in Experiment 3 (Fig. 1C, D), monkey OZ exhibited a bimodal SRT distribution, whereas monkey DZ exhibited a unimodal SRT distribution.

Upon bilateral FEF inactivation (blues traces in Fig. 1A and 1B and blue inverted histograms in Fig. 1C and 1D), SRTs increased by between ∼30 to 50 ms for saccades in either direction in both monkeys. A decrease in ES frequency paralleled this increase in SRT, with the proportion of ES decrease from 41 to 7% and 80 to 36% for rightward and leftward saccades for monkey DZ respectively, and from 41 to 32% and 59 to 42% for rightward and leftward saccades for monkey OZ respectively. Further, individual ES at even the shortest SRTs persisted when the FEF was inactivated (e.g., compare the short-SRT responses for monkey OZ when the FEF was inactivated or not). These results demonstrate the persistence of ES generation, even when a large volume of FEF was inactivated bilaterally.

### The effects of FEF inactivation on SRTs and express saccade frequency depend on target configuration

Rightward and leftward SRT distributions are represented as violin plots in Figs. 2 and 3 respectively, segregated by subject and by experimental condition. Recall that unilateral FEF inactivation was combined with potential targets located contralaterally in Exp1 or bilaterally in Exp2, and that bilateral FEF inactivation was combined with potential targets located bilaterally in Exp3 (i.e., there were no leftward saccades in Exp1). Each pair of red and blue violin plots represent the SRT distributions in the FEFwarm and FEFcool conditions, respectively; the numbers above the violin plots represent the median SRT, and the percentage of express saccades.

**Figure 2:**
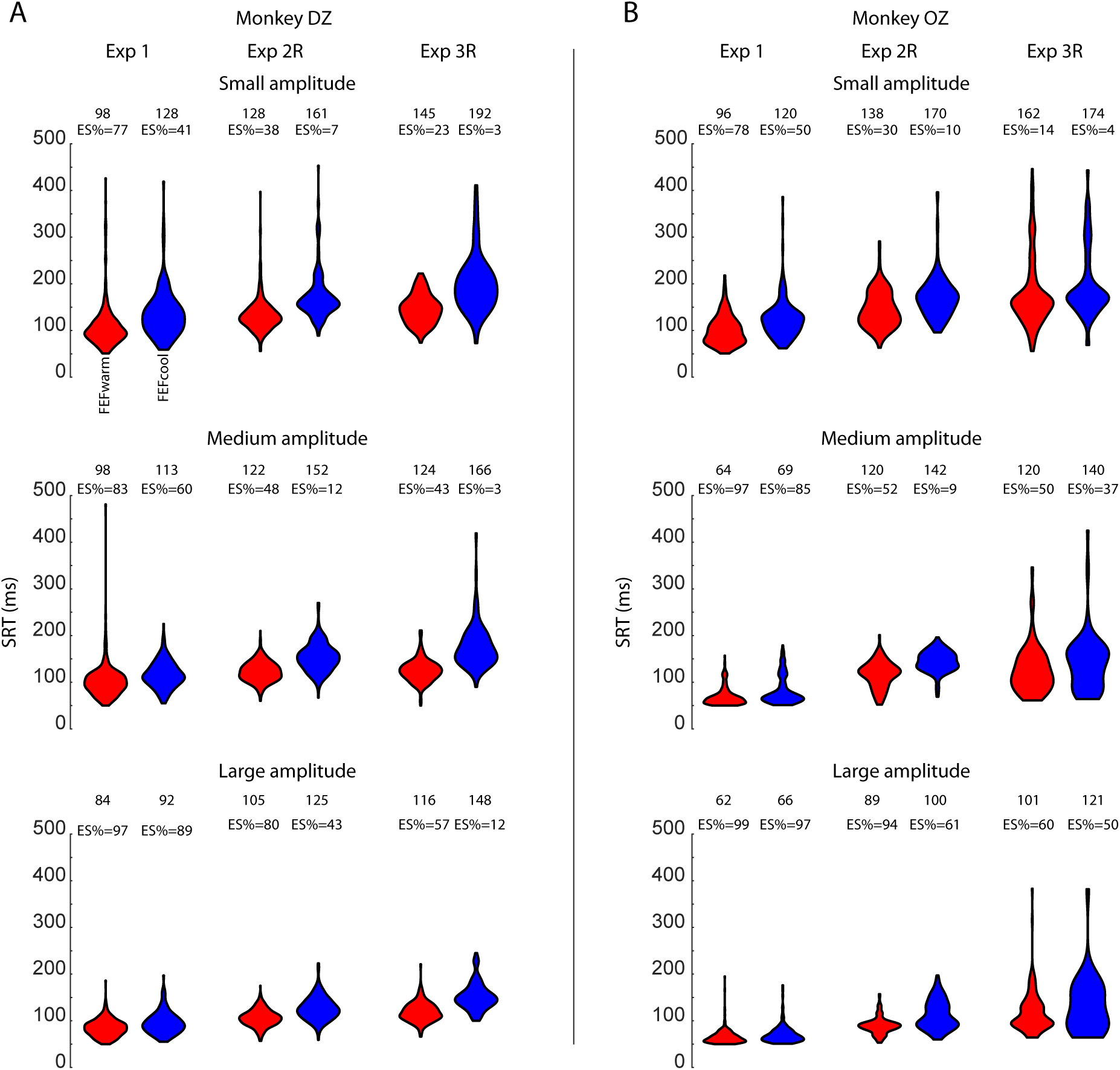
Effect of FEF inactivation on rightward SRT. SRTs for rightward saccades for monkey DZ (A) and monkey OZ (B). Within each plot, data are represented in violin plots depicting the distribution of SRTs. FEFwarm and FEFcool data is shown in red and blue, respectively. Each subplot shows data for Experiments 1 through 3, and different rows show data for different target amplitudes. The numbers associated with each violin plot provide the associated median SRT and ES frequency.

**Figure 3:**
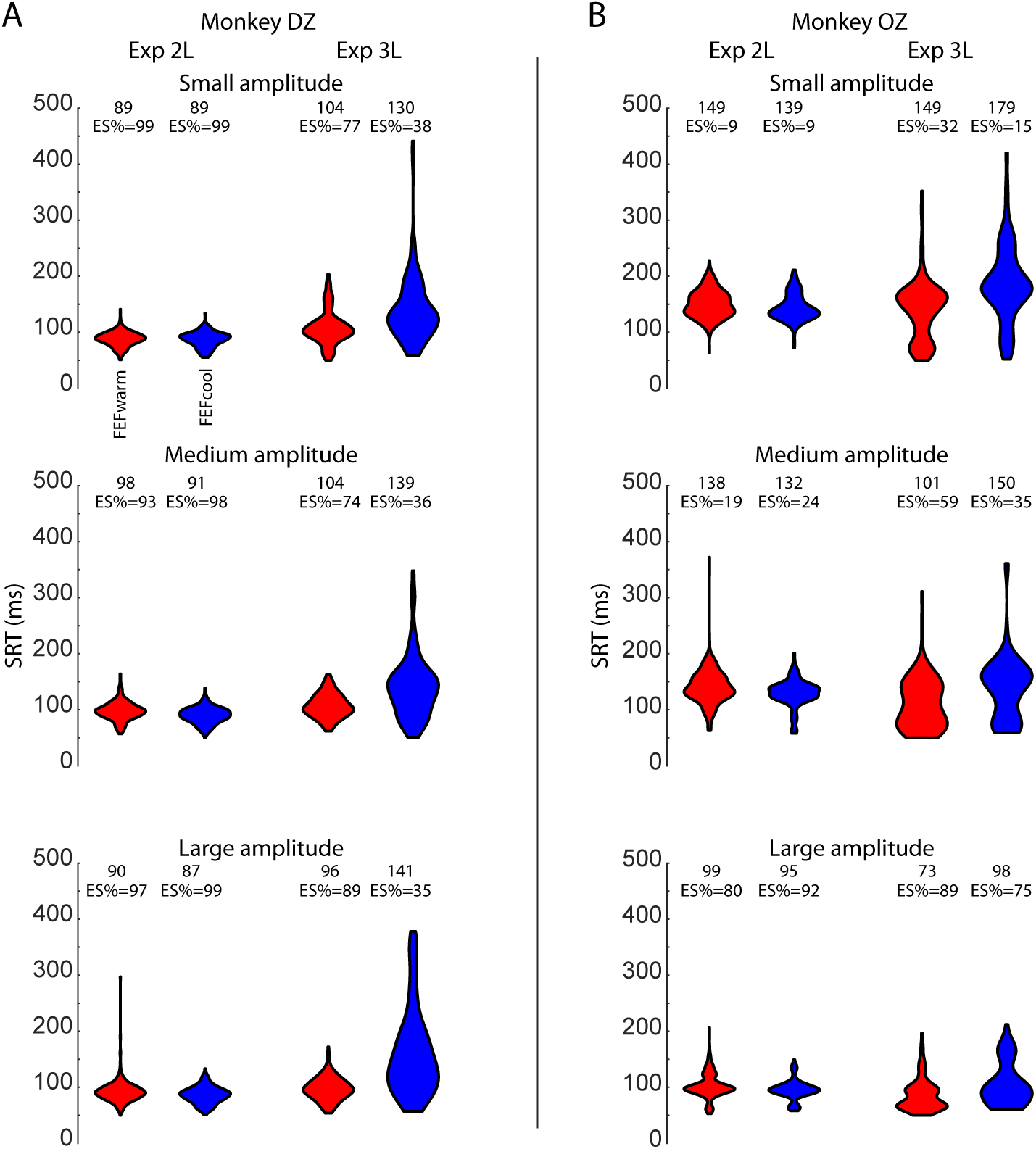
Effect of FEF inactivation on leftward SRT. SRTs for leftward saccades. Same format as for Fig. 2. Recall that there were no leftward targets in Experiment 1.

We quantified the effects of FEF inactivation on SRT by doing a 3-way ANOVA on each monkey with experimental configuration (Exp1, Exp2 and Exp3), amplitude (small, medium and large) and whether FEF was inactivated or not (FEFwarm and FEFcool) as 3 factors; separate 3-way ANOVAs were run for rightward and leftward saccades for each monkey followed by post-hoc tests corrected for multiple comparisons to establish the directionality of particular comparisons (all post-hoc tests were significant at p < 0.005, Tukey-kramer test). Speaking first to the SRTs of rightward saccades (Figure 2), we observed a significant main effect of inactivation (SRT increased with inactivation, Monkey DZ:F(1,4614)=868, p<0.005; Monkey OZ: F(1,3818)=194, p<0.005), amplitude (amplitude dependent shortening of SRT, Monkey DZ:F(2,4614)=356, p<0.005; Monkey OZ: F(2,3818)=596, p<0.005), and experimental configuration (SRT increases with increasing target uncertainty, Monkey DZ:F(2,4614)=616, p<0.005; Monkey OZ: F(2,3818)=890, p<0.005). We also observed significant interaction effects between inactivation and amplitude (bigger effect of inactivation on smaller amplitude saccades; Monkey DZ:F(2,4614)=28.13, p<0.005; Monkey OZ: F(2,3818)=3.34, p=0.03) and between inactivation and experimental configuration (bigger effect of inactivation with increasing target uncertainty and volume of inactivation (unilateral or bilateral; Monkey DZ:F(2,4614)=28.13, p<0.005; Monkey OZ: F(2,3818)=3.34, p=0.03).

Next, we report the effect of unilateral (left-FEF) or bilateral FEF inactivation on the SRT of leftward saccades. Previous work has shown that pharmacological inactivation of the FEF decreases ipsilesional SRTs (Dias and Segraves 1999; Sommer and Tehovnik 1997), whereas large volume cryogenic inactivation or radiosurgical ablation increases ipsilateral SRT (Kunimatsu et al. 2015; Peel et al. 2014). When the FEF was not inactivated, both monkeys generated a substantial proportion of leftward ESs (red violin plots in Fig. 3). Like rightward saccades we performed 3-way ANOVA with inactivation, amplitude and experimental configuration as factors and found a significant main effect of inactivation (Monkey DZ: F (1, 2967) = 291, p<0.005; Monkey OZ: F (1, 2036) = 47.5, p<0.005) and significant interaction effects between inactivation and amplitude (Monkey DZ: F (2, 2967) = 18.34, p<0.005; Monkey OZ: F (1, 2036) = 5.89, p<0.005) and between inactivation and experimental configuration (Monkey DZ: F (2, 2967) = 443.9, p<0.005; Monkey OZ: F (1, 2036) = 137, p<0.005). Posthoc analysis revealed a significant decrease in SRT for ipsilateral (leftward, Exp 2) saccades (Monkey DZ: p<0.005; Monkey OZ: p<0.005, Tukey-kramer test) while a significant increase in leftward SRT was observed during bilateral FEF inactivation (Exp 3; Monkey DZ: p<0.005; Monkey OZ: p<0.005, Tukey-kramer test).

### FEF inactivation decreases but does not abolish ESs

We next investigated the effects of unilateral-and bilateral FEF inactivation on ES frequency. The changes in ES frequency were largely predictable given the changes in SRT noted above, with greater decreases or increases in ES frequency accompanying larger increases or decreases in SRTs upon FEF inactivation. Accordingly, we will only emphasize the most important points in the analysis and present the highlights of an analysis of the changes in ES frequency conducted separately for each monkey for leftward and rightward saccades.

Without FEF inactivation, ES frequency depended on target configuration, being higher in Experiment 1 with only rightward potential targets (exceeding 75% in both monkeys; Fig. 2), than in Experiments 2 and 3 with bilateral potential targets. In both monkeys, ESs were generated most often to the most eccentric target locations. FEF inactivation almost always decreased ES frequency to a given target location, doing so to a greater degree in monkey DZ. Bilateral FEF inactivation also caused a greater decrease in ES frequency in monkey DZ than unilateral FEF inactivation, but this trend was not observed in monkey OZ. Bilateral FEF inactivation also decreased the prevalence of leftward ESs in Experiment 3, although unilateral inactivation of the left FEF increased the prevalence of leftward SRTs in Experiment 2 (Fig. 3). Regardless of these influences of FEF inactivation on ES frequency, it is important to stress that ESs were never completely abolished.

Finally, to complete our analysis of the changes in SRT and ES frequency with the various combinations of target and inactivation configurations, we specifically analyzed the SRTs of ESs made with or without the FEF inactivated. The range of SRTs classified as ESs in NHPs is quite broad (50-120 ms), hence it is unclear for example whether the decreases in rightward ES frequency is accompanied by an increase in the average SRT of rightward ESs, and whether the shortest SRT ESs continue to be generated upon FEF inactivation. In Figure 4, we present violin plots of SRTs specifically for ESs using the same format as Figs. 2 and 3. We performed a 3-way ANOVA analysis of rightward SRT as explained previously but limited SRTs to ES range. Similar to that seen for the full range of SRTs, we observed a main effect of both amplitude (Monkey DZ: F (2, 2474) = 11.3, p<0.005; Monkey OZ: F (2, 2410) = 38.92, p<0.005) and experimental configuration (Monkey DZ: F (2, 2474) = 151.28, p<0.005; Monkey OZ: F (2, 2410) = 218.56, p<0.005). Only monkey DZ showed a significant main effect of inactivation (Monkey DZ: F (1,2474) = 11.5, p<0.005; Monkey OZ: F (1, 2410) = 1.55, p=0.21). Both the monkeys did not show any significant interaction effects either between inactivation and amplitude or inactivation and experimental configuration (p>0.05 in all the cases). Posthoc analyses showed that rightward ES SRTs increased during unilateral FEF inactivation, but surprisingly, did not change during bilateral FEF inactivation (Monkey DZ: Exp1: p< 0.005, Exp2: p=0.01, Exp3: p=0.95, Tukey-kramer test; Monkey OZ: Exp1: p<0.005, Exp2: p=0.11, Exp3: p=0.41, Tukey-kramer test).

**Figure 4:**
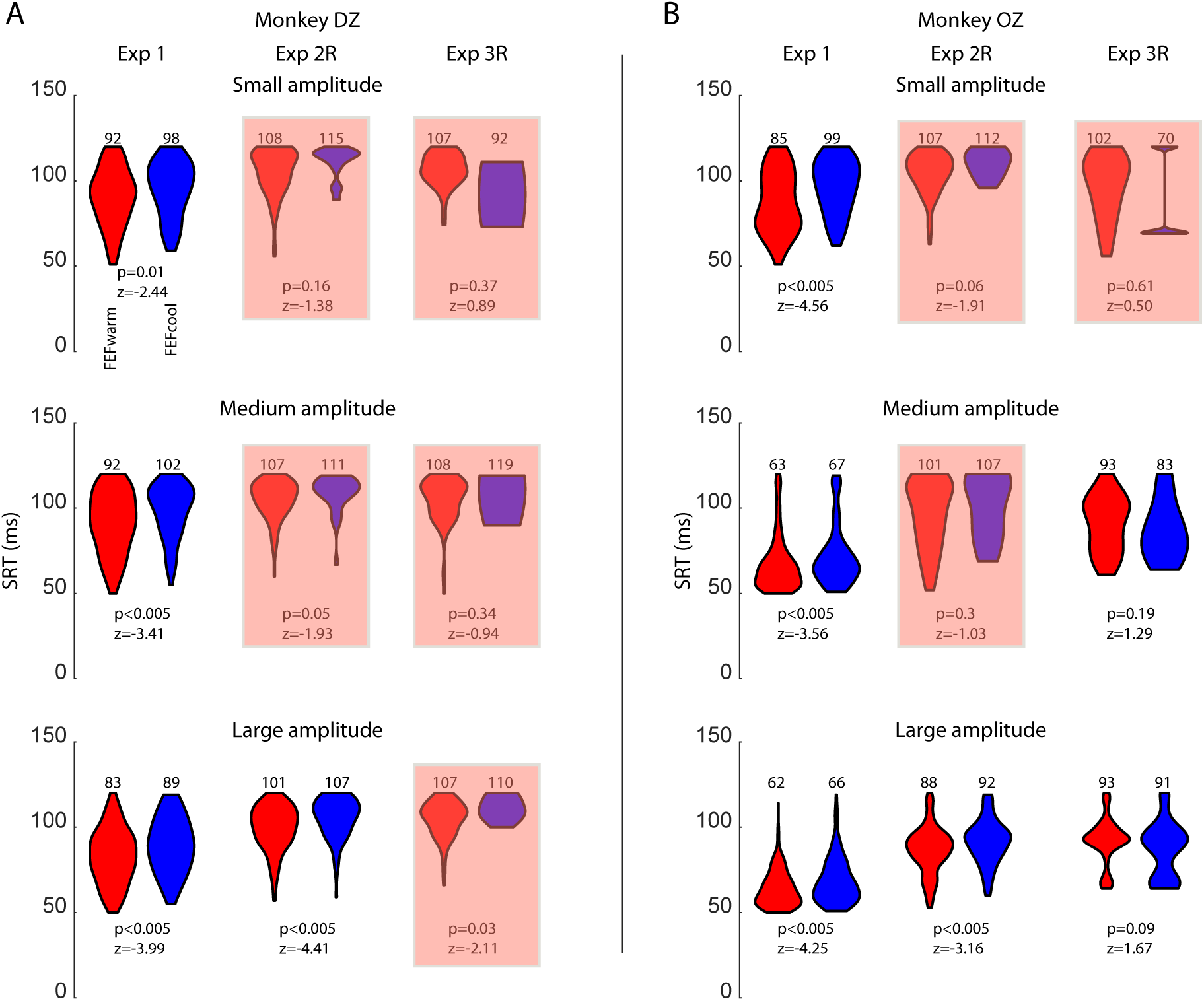
Effect of FEF inactivation on SRT of rightward ESs. SRTs for rightward ESs. Same format as Fig. 2, except that trials were only include if they fell in the ES range between 50-120 ms. Red shaded boxes represent configurations with an insufficient number of trials in the FEF cool condition to permit detailed analyses.

In the above analysis there were many subgroups with limited number of trials during the FEFcool condition (<20 trials, mostly belonging to small and medium amplitude saccades, shaded in red in Fig. 4). To check if a lack of datapoints were responsible for the preceding results we performed a 2-way ANOVA with experimental configuration and inactivation as two factors on data corresponding with large amplitude (20 deg) where enough data were available. Posthoc comparison of ES SRTs across experimental configurations revealed that SRT of rightward ESs increased following unilateral FEF inactivation (Exp 1 and Exp 2R) but did not change for rightward ESs during bilateral FEF inactivation in both the monkeys (Exp 3R; Figure 4, bottom row; monkey DZ: Exp1.: p<0.005, Exp2.: p<0.005, Exp3R.: p=0.63, monkey OZ: Exp1.: p<0.005, Exp2.: p=0.01, Exp3R.: p=0.36, Tukey-kramer test). We repeated the same analysis for leftward saccades and observed no change in ESs SRT following bilateral FEF inactivation in monkey DZ (data not shown; monkey DZ: p=0.97; monkey OZ: Exp3L: p<0.005, Tukey-kramer test). Further, although there were some situations where the shortest-SRT ESs were seemingly abolished to some targets (e.g., for the smallest amplitude targets in Experiment 1 for both monkeys), the general impression of these data is that both animals retained the ability to generate ESs at the shortest SRTs. Further, it is particularly impressive that FEF inactivation did not alter the modes of the bimodality apparent for ESs to large-amplitude targets in monkey OZ in Experiment 3 (lower-right plots in Fig. 4).

### FEF inactivation decreases accuracy of ESs

Pharmacological or cryogenic FEF inactivation is known to increase the targeting error and endpoint scatter of both visually-and memory-guided saccades, with larger deficits accompanying more complex tasks (Dias and Segraves 1999; Peel et al. 2014; Sommer and Tehovnik 1997). Consistent with the relative simplicity of the gap-saccade task, we observed only a modest increase in endpoint saccade during FEF inactivation, which is evident in the moderate increase in the dispersion of eye position traces during bilateral FEF inactivation in Experiment 3 (Fig. 1). Briefly, we performed a 3-way ANOVA to analyze changes in rightward saccade end point scatter with experimental configuration (Exp1, Exp2 and Exp3), amplitude (small, medium and large) and inactivation (FEFwarm and FEFcool) as 3 factors. We observed a significant main effect of experimental configuration (trend of increasing endpoint scatter from Exp1 through Exp3; Monkey DZ: F(2, 4614) = 38, p<0.005; Monkey OZ: F(2, 3818) = 102.8, p< 0.005) amplitude (increasing endpoint scatter with increasing amplitude; Monkey DZ: F(2,4614) = 30.9, p<0.005; Monkey OZ: F(2, 3818) = 91.5, p<0.005) and inactivation (increased endpoint scatter during FEFcool condition; Monkey DZ: F(1, 4614) =435.9, p<0.005; Monkey OZ: F(1, 3818) =27.39, p<0.005) in both the monkeys. We also observed a significant interaction effect between inactivation and experimental configuration, with larger increases in endpoint scatter upon FEF inactivation as we moved from Exp1 through Exp3 (Monkey DZ: F (2,4614) = 30.6, p<0.005; Monkey OZ: F (2, 3818) = 3.63, p=0.02). Posthoc analysis revealed significant increases in endpoint scatter in all three experimental configurations in monkey DZ (p<0.005, Tukey-kramer test), but only a tendency to increase in monkey OZ (Exp1: p=0.52, Exp2: p<0.005 and Exp3: p=0.12, Tukey-kramer test). Since previous studies have already established that endpoint scatter increases for regular latency saccades during FEF inactivation, we next specifically quantified the endpoint scatter of ESs. As mentioned in the previous section many groups did not have enough datapoints, so we focused on large amplitude trials (where we had enough trials) and conducted a 2-way ANOVA with experimental configuration and inactivation as factors. Only monkey DZ showed the main effect of inactivation for rightward saccades (p<0.005, 2-way ANOVA) with none of the individual group comparisons after correction for multiple comparison yielding a significant difference (monkey DZ: Exp1.: p=0.08, Exp2.: p=0.06, Exp3R.: p=0.72, Tukey-kramer test). Monkey OZ did not show significant results in any comparison. Further, endpoint scatter for ipsilateral saccades after unilateral FEF inactivation did not change in any of the subjects (p>0.05, 2-way ANOVA). Overall, the increase in endpoint scatter of ESs during FEF inactivation was quite modest, usually increasing by less than 1°.

Changes in targeting error with FEF inactivation mimicked the modest effects observed for endpoint scatter. We observed a significant increase in targeting error with changing experimental configuration (increased targeting error for Exp2 and 3 when compared to Exp1), amplitude (increased targeting error with increasing saccade eccentricity) and with inactivation (increased targeting error during FEF inactivation) in both the monkeys (p<0.005, main effects of 3-way ANOVA). Next, we looked specifically at targeting errors in ESs, concentrating only on large amplitude ESs. A 2-way ANOVA with experimental configuration and inactivation as factors revealed a non-significant trend for increasing targeting error during FEF inactivation in each experimental configuration in both the monkeys, but these did not reach significance after correcting for multiple comparisons (p>0.05, Tukey-kramer test). Thus, like endpoint scatter, targeting error increased marginally for ESs.

### FEF inactivation decreases peak velocity of ESs

Pharmacological or cryogenic FEF inactivation also decreases the peak velocity of contralaterally-directed saccades generated in a delayed visually guided or memory guided task (Dias and Segraves 1999; Peel et al. 2014; Sommer and Tehovnik 1997). We chose to analyse the effect of FEF inactivation on peak velocity in a different way, given that peak velocity is related to saccade amplitude via the main sequence relationship, and because of the moderate influence of FEF inactivation on saccade metrics. To address whether FEF inactivation influences the peak velocity of ESs, we conducted an analysis where we first selected pairs of trials, one each from the FEF warm and FEF cool condition, matched for the latency range (e.g., for either an ES or a regular-latency saccade) and closely-matched for saccade metrics (i.e., pairs had to have horizontal and vertical component amplitudes within 0.5 deg of visual angle). As shown in Fig. 5, we then plotted the peak velocities of each pair against each other, plotting the data from the FEF warm or FEF cool condition on the x- or y-axis respectively (the red or black data in Fig. 5 shows the comparison of ES or regular-latency saccades, respectively). The results of the statistical comparison of peak velocity are also presented in figure 5 (Wilcoxon sign rank test).

**Figure 5:**
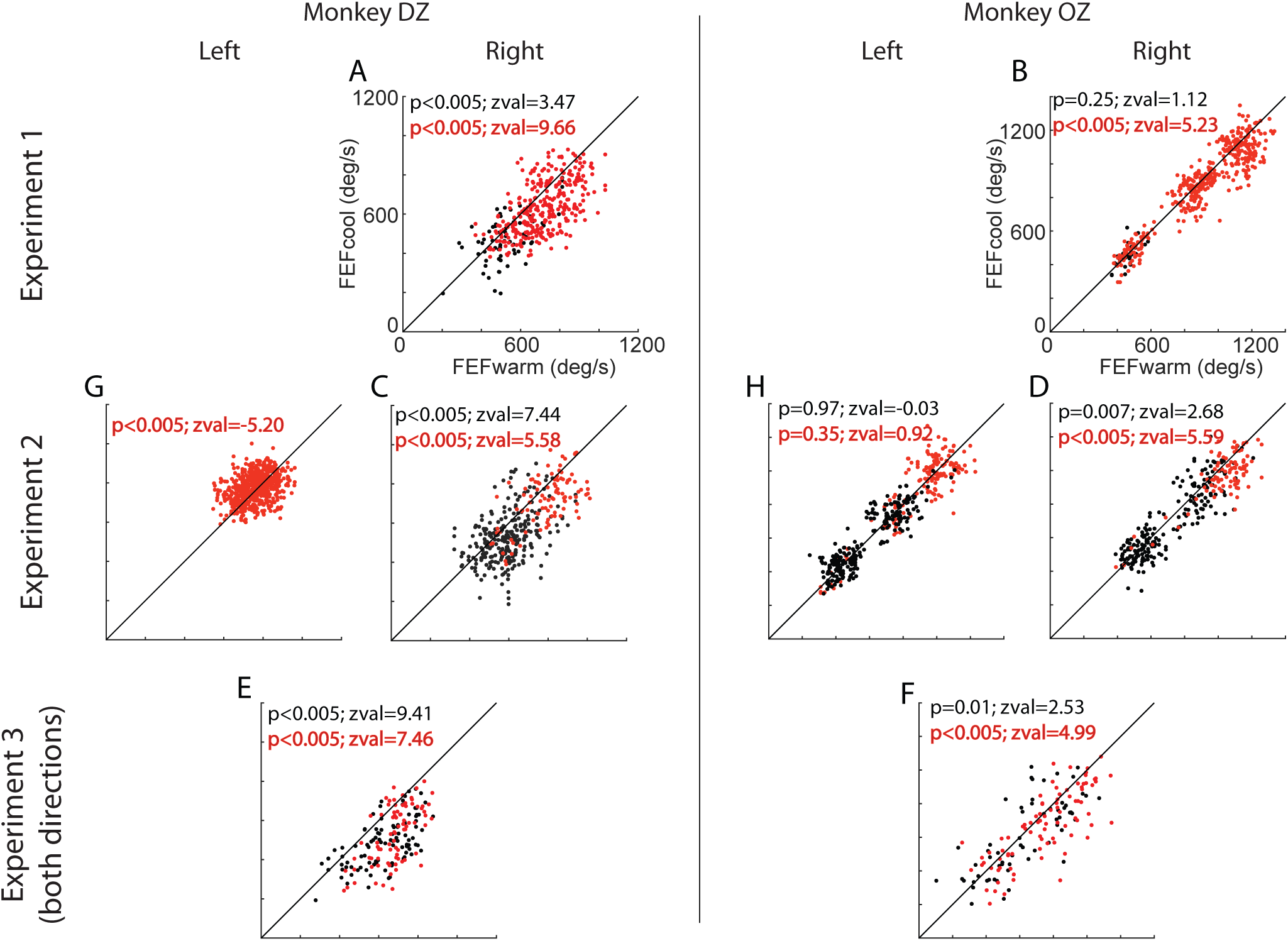
Effect of FEF inactivation on saccade peak velocity of regular latency saccades and ESs. A comparison of peak velocity of trials closely matched for metrics. Each point shows a pair of trials, collected either in the FEF warm condition (x-axis) or FEF cool condition (y-axis). Points are coloured by whether the saccade fell into the range of regular-latency saccades (black) or ESs (red). The results of statistical comparisons are given in the black or red text. The data from left and right saccades are pooled for Experiment 3, since the FEF was inactivated bilaterally.

The results of this comparison show that the peak velocity of both ESs and regular-latency saccades decrease during FEF inactivation. During Experiment 1 (Figure 5A & 5B) most of the selected pairs were in ES range (red) and showed a significant decrease in peak velocity. Very few trial pairs in regular latency range (black) could be extracted (especially in monkey OZ, figure 5B), but monkey DZ showed a significant decrease in peak velocity for both ES and regular-latency saccades. The median decrease in peak velocity for ESs and regular-latency saccades were 66 deg/s and 67 deg/s in monkey DZ and 21 deg/s and 1 deg/s in monkey OZ, respectively.

A similar result was observed in Experiment 2, although the decrease in peak velocity for contralateral (rightward) saccades appeared to be larger. Both the monkeys showed significant decreases in peak velocity for contralateral saccades (Figure 5D & 5F). The median decrease in peak velocity for ESs and regular-latency saccades for contralateral saccades were 94 deg/s and 69 deg/s in monkey DZ and 92 deg/s and 18 deg/s in monkey OZ, respectively. For Exp3 we pooled matched pairs from both directions and compared them and like Exp1 and Exp2 found decreased peak velocity after bilateral FEF inactivation (Figure 5G & 5H). The median decrease in peak velocity for ESs and regular saccades during Exp 3 were 134 deg/s and 107 deg/s in monkey DZ and 88 deg/s and 27 deg/s in monkey OZ, respectively. In summary, we observed a significant decrease in peak velocity of ESs during FEF inactivation, with the greatest decreases in Exp3.

Finally, we looked at the peak velocity of ipsilateral saccades (leftward saccades) during experiment 2. A previous study with unilateral FEF inactivation reported no changes in the peak velocity of ipsilateral saccades made during memory and delayed visually guided saccade tasks (Peel et al., 2014). In this study we found a similar result in one monkey (monkey OZ, Figure 5E) for both ESs and regular saccades, but observed that ipsilateral peak velocity increased during FEF inactivation in monkey DZ (median increase in peak velocity=15 deg/s; figure 5C). There were no trial pairs in regular saccade range in monkey DZ. In summary, even if we only consider the results from monkey DZ, the effect of unilateral FEF inactivation on ipsilateral peak velocity is modest.

## Discussion

To our knowledge, this is the first study examining the influence of large but reversible inactivation of unilateral or bilateral FEF on express saccades (ESs). We emphasize a number of novel results. First, even after bilateral FEF inactivation, ESs were not completely abolished despite decreasing in frequency. Second, the impact of FEF inactivation depended in part on the configuration of potential targets, being greater when potential targets were located bilaterally rather than unilaterally. Third, the peak velocity of ESs decreased during FEF inactivation. Together, these findings contribute to the understanding of the role of the FEF in the phenomenon of ESs, reaffirming the important but not critical role in setting up the conditions necessary for ES generation, and emphasizing that the influence of the FEF on saccade velocity regardless of SRT.

### Methodological considerations

In this study we inactivated the FEF using reversible cryogenic inactivation. This technique has a number of advantages compared to other temporary inactivation techniques or permanent ablation (Lomber et al. 1999), including a large volume of inactivation that impacts behaviour and neural activity within 2-3 minutes of changes in the temperature of the cooling loop. Such rapid onset and cessation of action, as well as the repeatability of inactivation within and across days, facilitates simultaneous electrophysiological recordings in interconnected areas (Dash et al. 2018; Peel et al. 2017). Another advantage of cryogenic inactivation is that it permits assessment of the functional consequences of loss of a large volume of tissue without confounds arising from neuroplasticity. The oculomotor performance of monkeys is surprisingly unimpaired after permanent ablation of the FEF following a period of recovery (Schiller et al. 1987; Schiller et al. 1980), and ESs are either unchanged or potentiated in human patients with frontal lesions (Braun et al. 1992; Guitton et al. 1985; Rivaud et al. 1994). Our results, by detailing the immediate decreases in ES frequency that accompany FEF inactivation, suggest that patients’ persistent capacity to generate ESs arises not from an inability to suppress reflexive glances consequent to frontal lobe damage, but instead to long-term functional recovery of oculomotor behaviours.

We cooled only the cryoloop placed in the inferior arm of the arcuate sulcus. Cooling this arm alone produced deficits ∼70% as large as the deficits observed when cooling both the inferior and superior arms together (Peel et al. 2014). Cyrogenic techniques inactivate tissue within a 1.5 mm radius of the cryoloop (Lomber et al. 1999), leading to an estimated volume of ∼90 mm^3^ of inactivated tissue in the anterior bank of the arcuate sulcus with a 7-8 mm length cryoloop (Peel et al. 2017). This volume is ∼3-6 times larger than the volume inactivation by pharmacological means, which range between 14-33mm^3^ (Dias and Segraves 1999; Peel et al. 2014; Sommer and Tehovnik 1997). Although it is difficult to ascertain directly, visual inspection of the tissue ablated in the studies by Schiller and colleagues (Schiller and Chou 1998; Schiller et al. 1987) and Kunimatsu and colleagues (Kunimatsu et al. 2015) show damage extending along the length of the arcuate sulcus and into the gyral crown, meaning that the total volumes of ablated tissue in these studies are perhaps ∼25-50% larger than the volume that we inactivated in the current study (see Peel et al. 2014 for further considerations). Our estimated volume of inactivation also only considers the volume lying on the anterior bank of the arcuate sulcus. We did not insulate the posterior aspect of our cryoloops, hence tissue lying on the posterior bank of the arcuate sulcus was inactivated as well. We have considered the potential impact of inactivation on the posterior bank more thoroughly in a previous publication (see Peel et al., 2014). Briefly, permanent (Rizzolatti et al. 1983) or reversible (Schieber 2000) lesions of tissue within the posterior bank of the arcuate does not produce the oculomotor deficits that we observed either in this or previous studies (Peel et al. 2014). Further, electrical stimulation of the posterior bank of the arcuate sulcus mainly evokes ipsilaterally-directed saccades (Neromyliotis and Moschovakis 2017) and saccade-related activity in the posterior bank is not biased contralaterally (Neromyliotis and Moschovakis 2018), in contrast to the FEF. We are therefore confident that the majority of the results reported here are attributable to the FEF within the anterior bank of the arcuate sulcus

Finally, each animal was implanted bilaterally. Although every attempt was made to ensure the symmetry of the implant relative to the spur of the arcuate, our results exhibit a fair degree of asymmetry both in the effects of inactivation, and in the SRT distributions and asymmetries within each animal even before inactivation. For example, the SRT distributions in monkey OZ but not DZ exhibited a clear bimodality, and monkey DZ tended to have shorter leftward than rightward SRTs. The effects of either unilateral or bilateral inactivation also tended to be larger in monkey DZ than OZ. Studies in NHPs involve a limited number of animals, and these types of idiosyncracies are unavoidable, and we place particular emphasis on results that were common in both animals.

### Comparison with other inactivation or ablation studies

Unilateral lesion or inactivation of the FEF produce a triad of contralateral saccade deficits, increasing SRT and decreasing peak velocity and accuracy (Acker et al. 2016; Dias et al. 1995; Dias and Segraves 1999; Kunimatsu et al. 2015; Lynch 1992; Peel et al. 2014; Schiller and Chou 2000; Schiller and Chou 1998; Schiller et al. 1987; Schiller et al. 1980; Sommer and Tehovnik 1997; van der Steen et al. 1986). Many of these studies examined saccade performance in the context of delayed or memory saccade paradigms in which ESs are not reliably generated. One exception is the 1987 ablation study by Schiller and colleagues, which found that contralateral ES frequency initially decreased when assessed 3 days after the lesion, but rapidly recovered back to pre-operative levels by 7 days post-lesion. Our findings show that the onset of FEF inactivation of contralesional ES frequency becomes apparent within minutes.

Recently, we reported that unilateral FEF inactivation decreases preparatory, low-frequency activity in the SCi during the gap interval (Dash et al. 2018). A component of this study found that ESs persisted during FEF inactivation, similar to what we found here. Importantly, this past study employed only two potential target locations, one in the center of the SCi neuron’s response field, and the other in the mirror opposite position. As we did not record SCi activity in the current study, we were free to use more configurations featuring more distributed target locations. The SRT distributions shown in Figs. 2 and 3 emphasize the influence of potential target configurations on the effects of FEF inactivation. Unilateral FEF inactivation caused a greater decrease in contralateral ES frequency and a corresponding larger increase in SRT in Experiment 2 (potential targets located bilaterally) compared to Experiment 1 (potential targets located unilaterally). This finding complements observations we and others have made on how the impact of FEF inactivation is greater for more complex oculomotor tasks (Dias and Segraves 1999; Peel et al. 2017; Peel et al. 2014; Sommer and Tehovnik 1997), extending this notion to immediate saccade tasks where the only difference is the number and location of potential targets.

The configuration of potential targets, and of task complexity in general, may also help resolve some confusion regarding the impact unilateral FEF inactivation on ipsilateral SRTs. Previous studies with unilateral FEF pharmacological inactivation have reported either negligible effects on ipsilateral saccades (Dias and Segraves 1999; Sommer and Tehovnik 1997) or an increase in premature ipsilateral saccades (Dias and Segraves 1999), in contrast to reports of increased ipsilateral SRTs following large cryogenic inactivation (Peel et al. 2017; Peel et al. 2014) or permanent ablation (Kunimatsu et al. 2015). Previously, we speculated that increases in ipsilateral SRT may be associated with larger volumes of inactivation (Peel et al. 2014), which could impact the contribution of sparsely-distributed saccade-related neurons tuned to ipsilateral directions (Crapse and Sommer 2009). In the current study however, we observed that ipsilateral SRTs can decrease during large-volume cryogenic inactivation (Fig. 3), resembling reports of decreased ipsilateral SRTs after permanent FEF ablation (Schiller and Chou 2000). An additional factor in determining the influence of unilateral FEF inactivation on ipsilateral SRTs, particularly in immediate saccade tasks, may be the number and distribution of potential target locations. In the work by Peel and colleagues (2014), targets could be presented at 1 of 32 locations distributed throughout oculomotor space, whereas the current study either 3 or 6 potential target locations distributed along the horizontal meridian.

### The FEF potentiates but is not critical for ES generation, and contributes to ES metrics and kinematics

The ablation work of Schiller and colleagues (1987) established that the integrity of the SC, but not FEF, is critical for ES generation. Consistent with this, we found using cryogenic inactivation that ESs are spared and can even be generated at the exact same SRT during either uni- or bilateral FEF inactivation. The FEF has direct projections to the SC (Kunzle et al. 1976; Stanton et al. 1988) that can relay signals that span the sensorimotor continuum (Everling and Munoz 2000; Sommer and Wurtz 2000) to influence SCi activity and oculomotor behaviour (Dash et al. 2018; Inoue et al. 2015; Peel et al. 2017). The FEF also projects directly to oculomotor regions downstream of the SC (Huerta et al. 1986; Leichnetz et al. 1984; Schnyder et al. 1985; Segraves 1992; Stanton et al. 1988), and this pathway presumably underlies the ability for monkeys to recover oculomotor abilities in the weeks to months after SC ablation. In the intact animal, however, it appears that the FEF predominantly influences saccadic behaviour through the SCi (Hanes and Wurtz 2001).

How do our results add to the understanding of the neural mechanisms preceding and during ESs? Our main result that ES frequency decreases but is not abolished by FEF inactivation is largely consistent with the idea that the FEF is one source of preparatory activity that help establish the preconditions that potentiate the generation of ESs. In a related experiment (Dash et al. 2018), we recorded SCi activity related to ESs either with or without unilateral FEF inactivation, and found that FEF inactivation reduced preparatory, pre-target information in the SCi. Such decreases in SCi preparatory activity related to both an increase in overall SRT and a decreased frequency of ESs during FEF inactivation. Surprisingly, SCi preparatory activity was largely unchanged when we specifically examined those rare instances where an ES was generated during FEF inactivation. Our interpretation, which holds during either unilateral or bilateral FEF inactivation, was that inputs from other cortical and subcortical areas, many of which also display increasing preparatory activity before target onset in related tasks (Ashmore and Sommer 2013; Chen et al. 2013; Hikosaka et al. 1989; Johnston et al. 2014; Kim and Lee 2017; Koval et al. 2011; Ohmae et al. 2017; Wurtz and Hikosaka 1986).

A novel finding in the current study is that inactivation of the FEF influences ES parameters, decreasing both accuracy and peak saccade velocity of ESs during either unilateral or bilateral FEF inactivation. ESs can be viewed as a form of a visual-grasp reflex, and the SCi does appear to be critical in the phenomenon of ESs. That being said, our finding that FEF inactivation decreases the accuracy and peak velocity of ESs are novel to our knowledge, and emphasizes that the subcortical mechanisms for ES execution do not operate in isolation from cortical inputs. In particular, the influence of FEF inactivation on saccade velocity was quite substantial, being of approximately the same magnitude as that for regular-latency saccades (Fig. 5). Correlates with saccade velocity have been found in saccade-related activity in the SCi but not the FEF (Reppert et al. 2018; Segraves and Park 1993; Smalianchuk et al. 2018; Waitzman et al. 1991). As mentioned above, decreases in contralateral saccade velocity are common following FEF inactivation, and our results emphasize that such velocity decreases persist for ESs. Whether this persistent influence on saccade velocity relates to a decrease in the vigor of saccade-related information in the SCi and/or to a loss of FEF input into the brainstem burst generator remains to be determined.

## Authors contributions

SD and BDC designed the experiments, SD conducted the experiments, SD and TP analyzed the data, SGL led the cryogenic surgeries, SD and BDC wrote the paper, SD, TP, SGL and BDC approved the manuscript.

## Acknowledgements

This work was supported by operating grants from the Canadian Institutes of Health Research to BDC (MOPs: 93796, 123247 and 142317) and the Natural Sciences and Engineering Research Council (NSERC; RGPIN-311680). SD and TRP were supported by funding from an NSERC CREATE grant and TRP was supported by an Ontario Graduate Scholarship. TRP is presently located in the Département de neurosciences, Université de Montréal, Montréal, QC, Canada

